# Simultaneous interneuron labeling reveals population-level interactions among parvalbumin, somatostatin, and pyramidal neurons in cortex

**DOI:** 10.1101/2023.01.09.523298

**Authors:** Christian Potter, Constanza Bassi, Caroline A. Runyan

**Author notes:** Contributed equally (Christian Potter, Constanza Bassi).

## Abstract

Cortical interneurons shape network activity in cell type-specific ways, and are also influenced by interactions with other cell types. These specific cell-type interactions are understudied, as transgenic labeling methods typically restrict labeling to one neuron type at a time. Although recent methods have enabled post-hoc identification of cell types, these are not available to many labs. Here, we present a method to distinguish between two red fluorophores *in vivo*, which allowed imaging of activity in somatostatin (SOM), parvalbumin (PV), and putative pyramidal neurons (PYR) in mouse association cortex. We compared population events of elevated activity and observed that the PYR network state corresponded to the ratio between mean SOM and PV neuron activity, demonstrating the importance of simultaneous labeling to explain dynamics. These results extend previous findings in sensory cortex, as activity became sparser and less correlated when the ratio between SOM and PV activity was high.

## Introduction

The spatiotemporal dynamics across thousands of excitatory and inhibitory synapses dictate how a neuron responds to a propagating signal^1^. Inhibitory interneurons (INs) can therefore have profound consequences on their local network, their activity altering spiking responses of individual neurons^2^ as well as network-level interactions across large populations of neurons^3^. The nature of these effects depends on an IN’s cell type, which is determined by its morphology, cellular connectivity, molecular expression, and firing properties^3^. A useful classification of interneuron subtypes is based on expression of parvalbumin (PV), somatostatin (SOM), or vasointestinal peptide (VIP), which divides interneurons into three non-overlapping groups. Although these are not homogeneous cell classes, neurons within them share many key functional properties, and the three types comprise almost all cortical interneurons^4^.

Of these three types, PV and SOM neurons most strongly inhibit pyramidal cells. PV neurons are preferentially targeted by feedforward excitatory inputs from outside the network, in parallel to pyramidal neurons that they also inhibit^5,6^, controlling their response gain^7,8^. SOM neurons more broadly integrate excitation from the local population^9^, and are more sensitive to modulatory input from other regions compared to PV^10,11^. The nature of these cells’ outputs is as distinct as their input patterns: PV neurons reduce activity by powerfully inhibiting the cell body, while SOM neurons inhibit the distal dendritic arbor, shaping how inputs are integrated by pyramidal neurons^4,12^. SOM neurons additionally inhibit PV neurons^13^, meaning that their activity can shift the mode of inhibition a pyramidal neuron receives. Experimental and computational work has suggested that this shift in activity corresponds to differences in cortical coding: changing the dynamic range of input a neuron receives^12^, and causing activity to become sparser^14^ or less variable from trial to trial^2^. To study the interplay between PV and SOM neurons, and the impacts of their dynamics on the local population, these neurons need to be simultaneously monitored.

To characterize the function of specific cell classes using fluorescence microscopy, experimenters often non-specifically express a green calcium indicator in all neurons and use genetic tools, such as Cre/lox recombination to selectively express a red fluorophore in one cell type of interest^15–18^. This approach is thus limited to monitoring the activity of one labeled cell type along with other neurons in the population, and does not allow the observation of interactions between specific cell types and the broader activity of the population. As a result, the ability to look at dynamics on individual trials is limited. A more powerful approach would allow the simultaneous recording of multiple cell types. Tools do exist to molecularly classify many cell types in tissue slices after *in vivo* imaging has been completed, but this process involves perfusing the animal, slicing the brain, staining for cell-type specific proteins, and registering the stained tissue to the *in vivo* images^19–22^. Although an established method, it is a technically challenging process that is not accessible to most labs.

To test the idea that the relative level of PV and SOM neuron activity corresponds to distinct population activity regimes, we used Cre- and Flp-dependent recombination to express distinguishable red fluorophores in PV and SOM neurons in the same mice. We exploited the differing two-photon excitation spectra of tdTomato and mCherry^23^, discriminating PV/tdTomato and SOM/mCherry neurons from each other by their relative fluorescence intensity across a range of excitation wavelengths from 780 – 1100 nm. This allowed us to identify the two cell types using red wavelength emission, freeing the green emission channel to measure GCaMP fluorescence changes, monitoring the activity of the full neuronal population. We observed highly separable population activity events characterized by bursts in activity of one neuron type but not the other. PV events corresponded to greater levels of shared variability across more densely active putative pyramidal neurons, while SOM events corresponded to sparser, decorrelated activity across the putative pyramidal population. Our method, which can be applied with any standard two-photon microscope with a tunable laser and two detection channels, has revealed how specific interactions between cell types affect the broader population, which would not have been possible if labeling only one cell type per animal.

## Results

### mCherry and tdTomato can be reliably identified by differences in their excitation spectra

TdTomato and mCherry are red fluorophores that are not easily distinguished by their emission spectra. However, their two-photon excitation spectra differ markedly, so in principle mCherry and tdTomato-expressing neurons could be distinguished by comparing their fluorescence intensity across excitation wavelengths^23^. To test for this possibility *in vivo*, we injected a Cre-dependent tdTomato construct (AAV1-pCAG-FLEX-tdTomato-WPRE) and a Flp-dependent mCherry construct (AAV1-Ef1a-fDIO-mCherry) in adjacent, but physically separate, locations in layer 2/3 of the posterior parietal cortex (PPC) of SOM-Flp (+/-) x PV-Cre (+/-) mice (Figure 1A). Next, we imaged the two sites within the same field of view *in vivo* across a range of excitation wavelengths. Even by eye, mCherry and tdTomato were distinguishable at different wavelengths, as the mCherry but not tdTomato site was visible at 800 nm excitation (Figures 1B and S1). We took the mean intensity of the pixels in each red cell body’s region of interest (ROI) at evenly spaced wavelengths every 20 nm from 780 to 1100 nm (excluding 1020 and 1060 nm, see STAR Methods), revealing separable excitation spectra between the neurons in the adjacent regions (Figures 1B and 1C).

**Figure 1.**
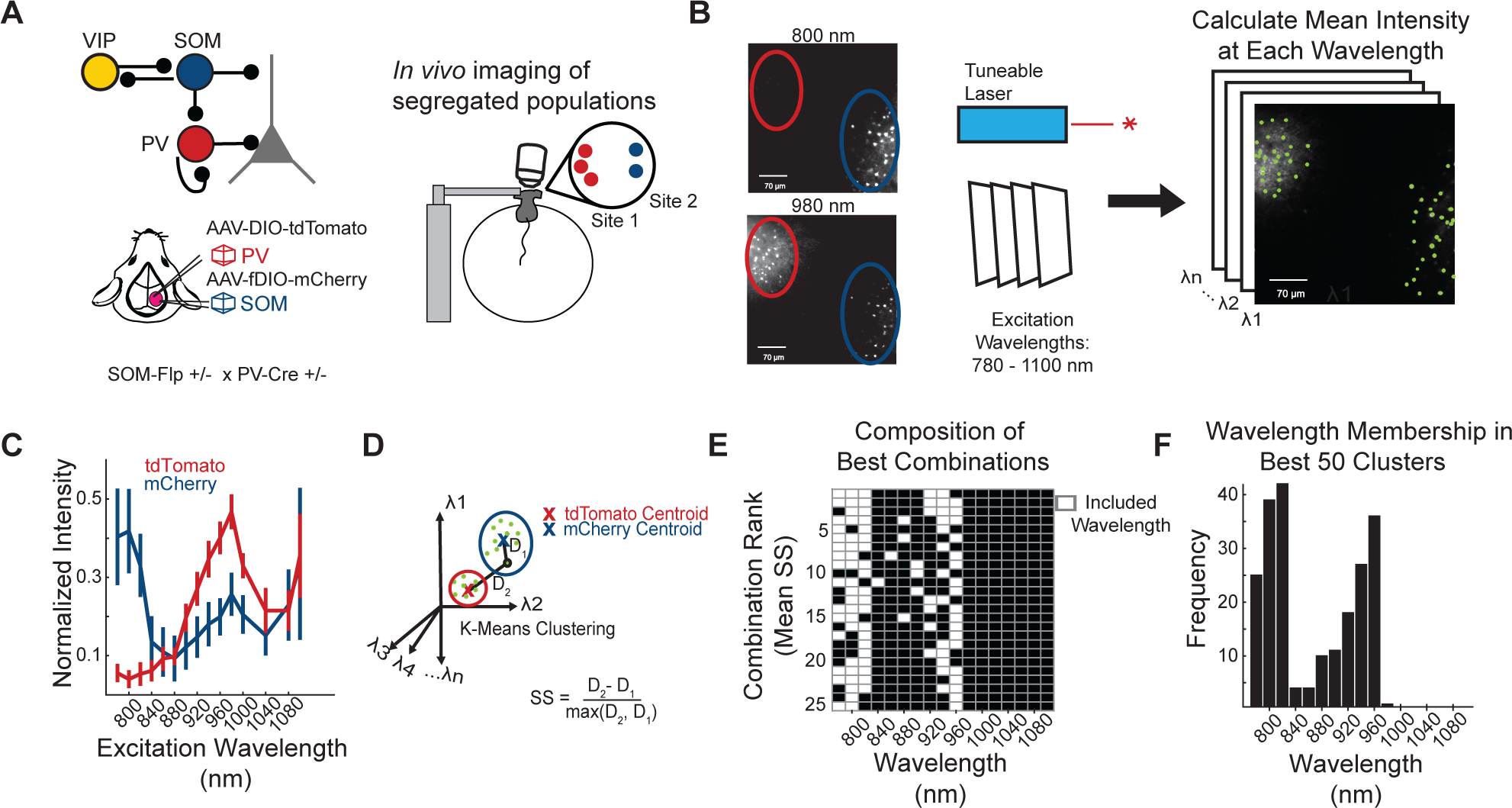
Validation of method to spectrally separate mCherry and tdTomato expressed in PV and SOM neurons *in vivo*. (A) SOM-Flp x PV-Cre mice received two adjacent, but physically segregated, injections of a Cre-dependent tdTomato and a Flp-dependent mCherry construct. For *in vivo* two-photon imaging, mice were head fixed on a spherical treadmill. (B) A field of view was selected to include both the tdTomato (red circle) and mCherry (blue circle) injection sites. A series of images was collected, with excitation wavelengths spanning from 780 to 1100 nm. The mCherry site, but not the tdTomato site, was visible with 800 nm light (left), while both sites were visible with 980 nm excitation. ROIs were drawn over the cell bodies of the neurons to select pixels for analysis, indicated by green masks. (C) Mean normalized intensity of tdTomato ROIs (red) and mCherry ROIs (blue), at all excitation wavelengths. Error bars indicate one standard deviation from the mean. The 1020 and 1060 nm wavelengths do not have a value since they were not used (see STAR Methods) (D) Schematic of the clustering strategy to identify tdTomato+ and mCherry+ neurons, based on their excitation spectra. K-means clustering was performed on the vectors of mean intensity values for all ROIs, specifying that they should be clustered into two categories. (E) K-means clustering was applied to the fluorescence of all mCherry and tdTomato ROIs collected at all possible combinations of two or more excitation wavelengths. For each combination, cluster separability was assessed by computing the mean of all silhouette scores, which is a measure of each ROI’s distance to the centroids of the two clusters. The 25 best combinations of wavelengths (white rectangles) for cluster separability are shown here, ranked by their mean silhouette score. (F) Frequency of a given wavelength occurring in the top 50 ranked combinations of wavelengths. 2 mice were used for panels C-F, with 6 different imaging sites across these animals, n = 326 ROIs.

To sort the ROIs into tdTomato+ and mCherry+ groups, we used k-means clustering for classification, as it is a simple algorithm that allowed us to specify two clusters for our data and quantify the separability of classification for each ROI (Figure 1D). Given that the separability of the two fluorophores varied across excitation wavelengths (Figure 1C), we hypothesized that a small subset of ideal excitation wavelengths would separate the ROIs more accurately than using all data. By determining which wavelengths produced the most separable clusters, we could reduce the number of neurons that would have to be excluded from analysis due to poor clustering performance. To find the highest performing excitation wavelength subsets, we performed k-means clustering on every possible combination of wavelengths and recorded the separability of each ROI, which is proportional to the confidence in classification. ROI separability was quantified by computing the silhouette score (SS), which is equal to the distance to the centroid of its assigned cluster subtracted from the distance to the other cluster’s centroid (Figures S2A and 1D). The mean SS of all ROIs for each combination was strongly related to classification performance, and most combinations of wavelengths achieved >95% classification accuracy (Figures S2B and S2C). To optimize classification performance before applying it to data with intermixed mCherry and tdTomato expression, where the data could become noisier due to neuropil contamination, we ranked the combinations of wavelengths by their mean SS scores (Figure S2D). This revealed certain wavelength combinations that produced more separable clusters. Unsurprisingly, these combinations tended to include wavelengths with greater differences in fluorescence intensity in the tdTomato- and mCherry-expression sites (Figures 1C, 1E, and 1F).

To define a threshold for cluster separation in each individual neuron, we examined the distribution of SSs from incorrectly classified neurons using ‘suboptimal’ wavelength combinations, the combinations that included excitation wavelengths with more similar fluorescence intensity in mCherry and tdTomato sites. This allowed us to simulate clustering on noisy data (Figure S2E). We chose a conservative exclusion threshold for the SS of each neuron such that 98% of misclassified neurons would be excluded from analysis, even in this unrealistically noisy data where mCherry and tdTomato were more poorly separated overall. Comparison to SSs from clustering on our intermixed expression sites (0.93 ± 0.13, mean ± s.d.) demonstrates how much noisier our simulated data was (0.85 ± 0.20, p = 7.64 * 10^-111^, Wilcoxon rank-sum), and still this conservative estimate resulted in the exclusion of only 6.7% of neurons in intermingled data in PPC (Figure S2F). Having performed this control analysis, we could be confident that our labeling approach would rarely yield misclassified neurons after excluding neurons with worse cluster separation.

### Simultaneous imaging of SOM, PV, and PYR neurons in layer 2/3 to study population-level interactions across cell types

Having validated our method for spectral separation of mCherry and tdTomato *in vivo* using physically separated control expression sites (Figure 1), we applied our approach to identify intermingled somatostatin-expressing neurons that were mCherry+ (SOM), parvalbumin neurons that were tdTomato+ (PV), and putative pyramidal (PYR) neurons that were negative for both red fluorophores in layer 2/3 of PPC. Given that interneurons comprise only ∼10-15% of neurons in the rodent cortex, and SOM and PV neurons make up ∼70% of these, we would expect ∼95% of the remaining neurons to be pyramidal^4^. All three neuron types expressed GCaMP6f, a genetically encoded calcium indicator to monitor and relate their ongoing spike-related activity^24^ (7 mice, 12 imaging fields of view). During imaging sessions, mice were head-fixed under a two-photon microscope, and voluntarily ran on a spherical treadmill (Figure 1A). In a typical field of view, 39±4 PV, 14±7 SOM, and 323±145 PYR neurons could be identified (mean ± s.d, Figures 2A and 2D).

**Figure 2.**
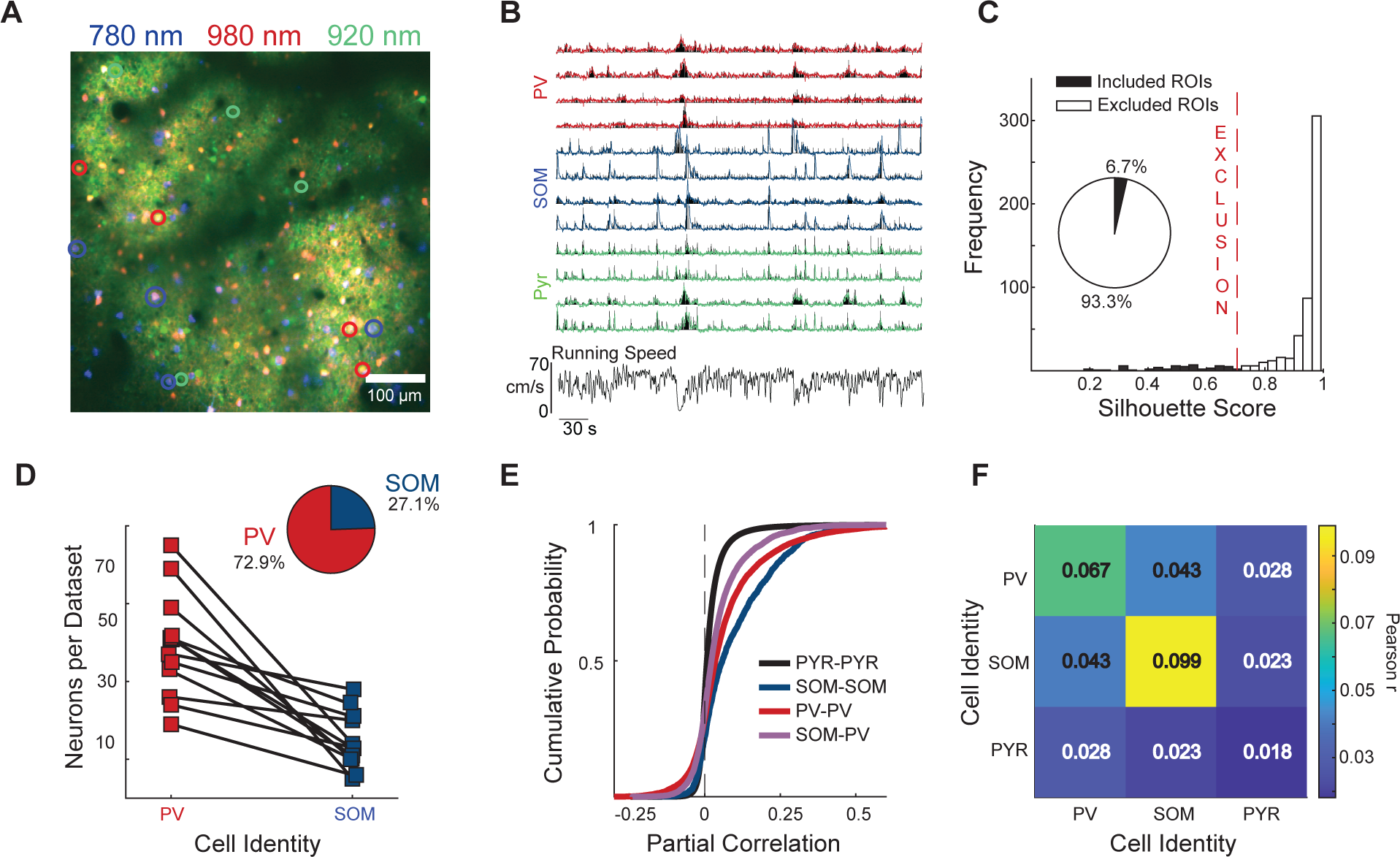
The strongest pairwise correlations are within interneuron subtypes. (A) Example field of view from layer 2/3 of PPC. The image is a merge of red fluorescence collected using 780 nm excitation (blue, visualizes mCherry+/SOM+ neurons), 980 nm (red, visualizes both mCherry+/SOM+ and tdTomato+/PV+ neurons), and green fluorescence collected using 920 nm excitation (green, visualizes all neurons). Neurons with green but not red fluorescence were considered putative pyramidal neurons (PYR). Colored circles indicate cells used for the traces in (B). (B) GCaMP fluorescence changes in a subset of the PV, SOM, and PYR neurons in the field of view in A (colored traces: red/PV, blue/SOM, and green/PYR). Black: estimated spike rates. Bottom: Time-aligned running speed of the mouse. (C) Distribution of tdTomato/mCherry cluster separability (silhouette scores) for all red neurons. 0.7 was selected as an exclusion threshold, based on our estimate of our simulated noisy data (see Figure S2). (D) Number of neurons classified as PV+/tdTomato+ and SOM+/mCherry+ in all datasets. (E) Cumulative distribution function of all pairwise correlations between neurons across the entire imaging session. Pairwise correlations were computed as partial correlations, regressing off any shared activity due to running. (F) Matrix showing the mean partial correlation within and between cell types. The mean correlation was significantly different between all non-symmetrical entries of the matrix (See Table S1 for more detailed statistics). Data is from 12 imaging sessions in 7 mice. Our labeling method yielded an average of 39±4 PV, 14±7 SOM, and 323±145 PYR neurons per dataset (mean ± s.d). Correlation results included 573661 pairs of PYR-PYR neurons, 1471 pairs of SOM-SOM, 10021 pairs of PV-PV, 6720 pairs of SOM-PV, 53661 pairs of PYR-SOM, and 135258 pairs of PYR-PV.

The activity of inhibitory interneurons is strongly correlated with the activity of other inhibitory interneurons of the same subtype, across cortex^17,22,25–27^. To determine whether these patterns of activity relationships between pairs of individual SOM, PV, and PYR neurons hold in mouse PPC, we computed Pearson correlations between each neuron pair across the imaging session, after discounting the effects of running velocity on each neuron’s activity (see STAR Methods). The distribution of pairwise correlations among cell type pairs was consistent with those previously observed in sensory cortex, with positive correlations among inhibitory neurons^22,25,27^, and SOM-SOM pairs were the most strongly and positively correlated (Figures 2E and 2F). Of all the cell type combinations, the least correlated pairs were the PYR-PYR pairs (Table S1).

### Identifying periods of differential interneuron activity

Interneuron subtypes tended to be positively correlated with other neurons of the same type (Figures 2E and 2F). Strikingly, when visually inspecting the activity patterns in the simultaneously imaged PV and SOM neurons, we also observed spontaneous events of elevated firing across neurons that were restricted to one cell type or the other (Figures 2B and 3A). We hypothesized that PV and SOM activity events relate to different activity states in the PYR population.

**Figure 3.**
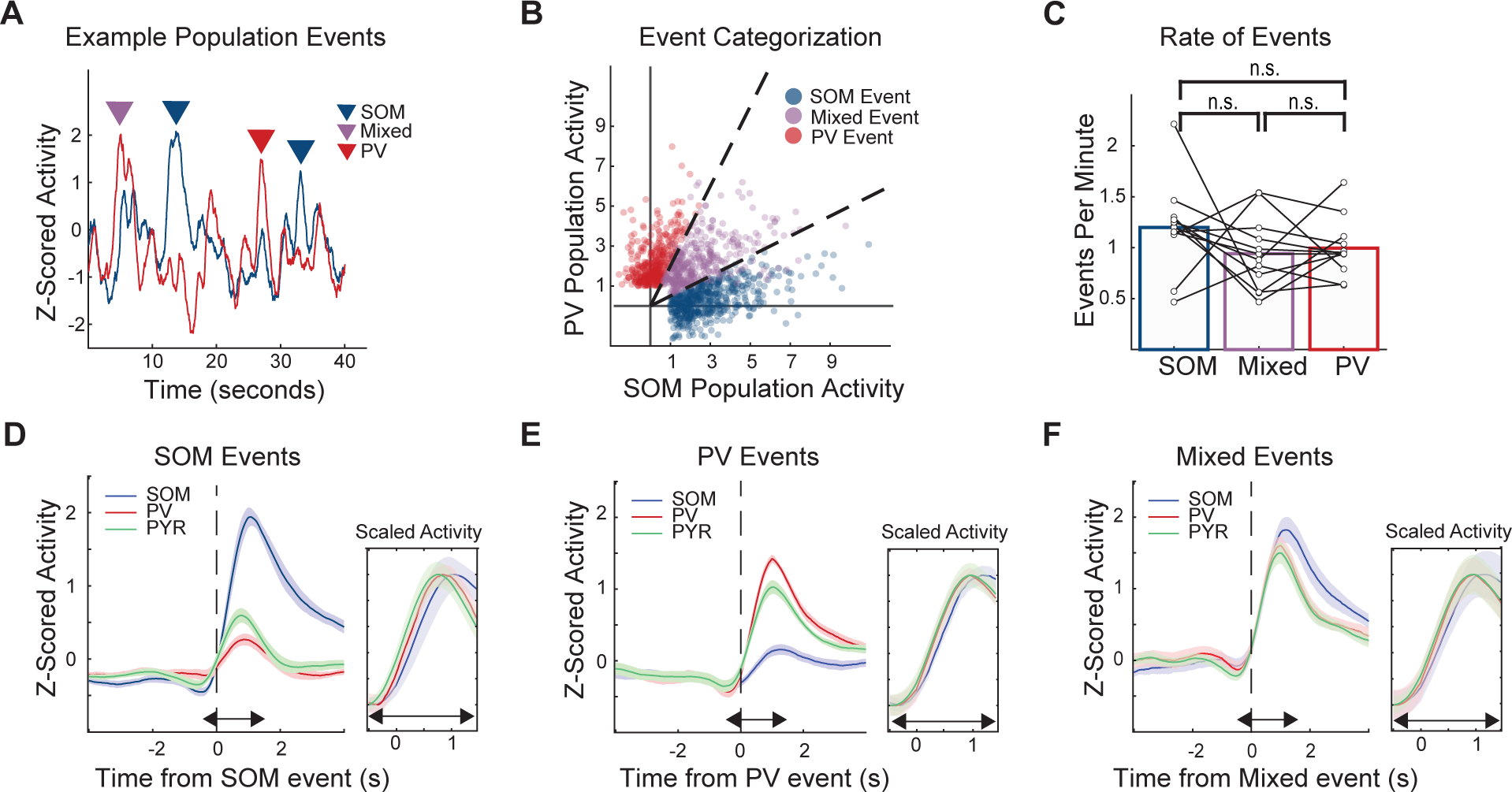
Interneuron event identification and timescale of mean activity during events. (A) Normalized population activity in simultaneously imaged SOM (blue) and PV (red) neurons (z-scored and averaged across neurons). Coordinated activity events were defined as peaks of activity that were unique to one population. Arrows above the activity traces mark these peaks. (B) SOM population activity plotted against PV population activity during events across all datasets (red: PV events, blue: SOM events, purple: Mixed). (C) Rate of events per minute. There was no significant difference between event types. (D) Activity of the SOM, PV, and PYR neurons aligned to the onset of SOM events. The mean activity for each cell type is z-scored across the entire session for each dataset, and then averaged across all events for each cell type. (E,F) As in D, but for PV and Mixed events. For all panels, “n.s.” means non-significant. Data comes from 12 imaging sessions in 7 animals, which yielded 430 SOM events, 400 PV events, and 387 Mixed events across all sessions.

First, to identify PV and SOM activity events, we used a peak finding algorithm to identify periods of elevated PV or SOM population activity and measured the mean activity of the other cell type population during this period (Figure 3A; see STAR Methods). Because elevated activity periods of one IN subtype could occur either independently or concurrently with activity in the other subtype, we defined three types of events, based on the relative activity of SOM and PV. For SOM and PV events, the normalized activity of one cell type was more than double the opposing type. For “Mixed” events, the ratio between the types of activity was less than two (Figure 3B). SOM events, PV events, and “Mixed” events were not statistically different in their frequency or duration across categories (Figure 3C; see Table S2 for statistics).

PPC is involved in navigational decisions, and neural activity in PPC can be sensitive to the animal’s running speed and direction^17,28^. While the mice in this study were not performing a task, they were running voluntarily on a spherical treadmill during imaging sessions. To determine whether PV, SOM, and Mixed activity events corresponded to behavioral events, we examined the running patterns of the mice. On average across mice, IN events corresponded to a small change in running acceleration; however, the mean peak of acceleration was not significantly different between event types and was not evident in all imaging sessions (p = 1.0, Kruskal-Wallis; Figure S4). Next, to relate IN activity events to overall population activity levels, we examined the activity of the PYR population during SOM, PV, and Mixed events. PYR activity was elevated during all three types of IN events; however mean PYR activity was significantly higher during PV and Mixed events compared to SOM events (dF/F mean ± s.d.: 0.21 ± 0.09 SOM, 0.23 ± 0.09 PV, 0.24 ± 0.09 Mixed; SOM/Mixed and SOM/PV p = 0.001, Wilcoxon Sign-Rank) but was not statistically different between PV and Mixed events (p = 0.52; Figures 3E and 3F). We plotted the time-course of mean activity scaled to its peak for each event, revealing that the peak activity of SOM neurons, which is likely the result of pooled input from local pyramidal neurons^4^, lagged mean PV activity by ∼200 ms during SOM (mean lag: 0.24 seconds, p = 0.042, Wilcoxon signed-rank) and Mixed events (Mean lag 0.17 seconds, p = 0.012), reflecting previously observed dynamics in V1^2,29^. In contrast, SOM activity did not lag behind PV activity during PV events (mean lag: 0.03 seconds, p = 0.50), potentially reflecting its reduced activity relative to PV.

### Comparing network states during differential periods of IN activity

When PV and SOM neurons are optogenetically stimulated as populations, they have different effects on local population activity dynamics^14,30^ and in shaping the response properties of individual neurons^2,8,9,31^. So far, our results suggest that PV and SOM neurons can act as highly coordinated and independent subpopulations. In the current study, we also have the unique opportunity to determine the changes in PYR network activity state that accompany these <u>natural</u> bursts in PV, SOM, or PV & SOM population activity.

We examined the activity of individual PYR neurons during IN events, by binning their activity from the peak of the event until two seconds after the IN activity peak. This accounted for most of the period of elevated IN activity across events (Figure S3D; Table S2). Consistent with the preferential connectivity and activity relationships between some pyramidal neurons and specific inhibitory cell types^25,32^, we did find small subsets of PYR neurons that tended to be more active either during SOM or PV events, as measured by differences in their mean activity for each event type over the analyzed period (Figure 4A). Although the majority of PYR neurons (80%) had no response preference, 17% of PYR neurons had significantly higher firing during PV events and 4% of neurons significantly higher firing during SOM events (Figure S5A; significance determined by p < 0.05 from a permutation test between firing rates for SOM and PV events). Activity of SOM-preferring neurons was significantly less correlated with the population than activity of PV-preferring neurons (p < 6.14*10^-8^, Wilcoxon rank-sum) or of neurons without preference (p = 0.016) over periods of non-event activity (Figure S5A). SOM-preferring and PV-preferring neurons were correlated with neurons with similar preferences across datasets (p = 0.91, Wilcoxon signed-rank), suggesting that these PYR subsets may form functional subnetworks that are relevant outside of IN events (Figure S5B).

**Figure 4.**
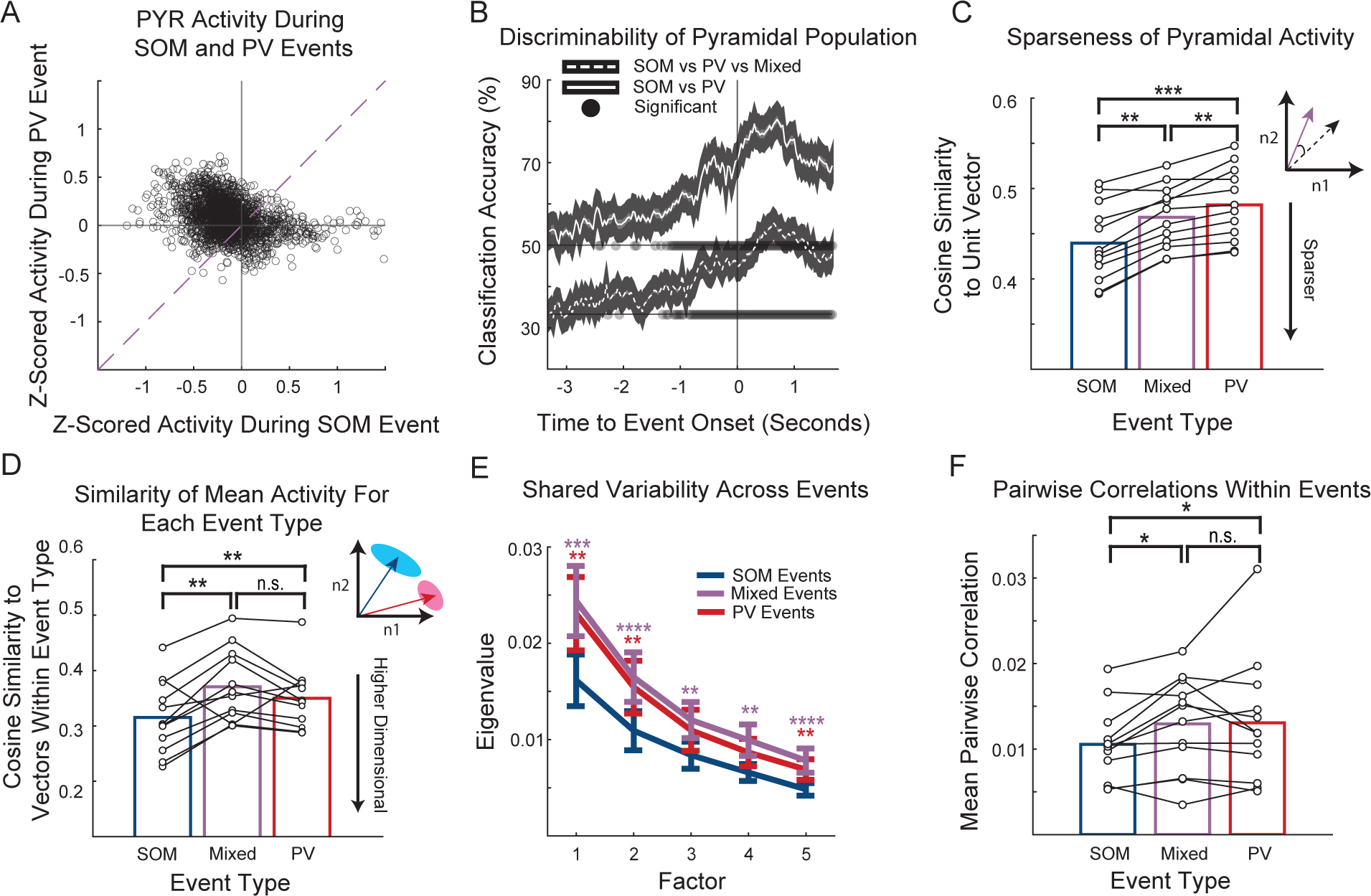
PV, SOM, and Mixed events are associated with different pyramidal population activity states. (A) Mean normalized activity for each pyramidal neuron during PV and SOM events. The activity of pyramidal neurons for each event was z-scored across all PV and SOM events for each neuron, and the mean of this z-scored activity was plotted. The unity line would indicate equal activity for SOM and PV events. (B) Performance of decoders trained to distinguish between SOM and PV events, or SOM, PV and Mixed events based on the mean activity during 300 ms bins. Shaded circles indicate classification accuracy that was significantly (p < 0.05) greater than the performance of a classifier with randomly shuffled labels for training, after Bonferonni correction for multiple comparisons at all 150 timepoints. (C) Sparseness of pyramidal neuron activity, measured by the mean angle between all vectors for an event type and the unit vector, which corresponds to all neurons being equivalently active. Vectors included each neuron’s activity, binned over two seconds following the event peak. (D) Pairwise cosine similarity of mean activity vectors 2s after the peak of the event. Pairwise similarity of pyramidal activity during SOM events was significantly lower than during PV events, indicating population activity spanned a wider space of possible activity patterns during SOM events. (E) Factor analysis was able to explain more shared variability in pyramidal neurons during PV and Mixed events compared to SOM events. Purple stars indicate significance between SOM and Mixed events, while red stars indicate significance between SOM and PV events. Factor analysis was performed 5 times on each dataset, subsampling 75 pyramidal neurons each time. (F) Pairwise correlations calculated between pyramidal neurons during the 2 seconds after the peak for each event. The mean pairwise correlation was significantly higher during PV and Mixed events compared to SOM events across datasets. For all panels, n.s. means no significance; *, p<0.05; **, p<0.01; ***, p<0.001, ****, p< 0.0001.

Previous work that has shown that functionally relevant subnetworks of Ins can be identified based on how they are driven by pyramidal activity^32^. Given our result of pyramidal preference for a particular IN event type, we hypothesized that PV and SOM events could be predicted by the pattern of PYR activity, even before the IN event onset. We aligned neural activity to IN event onsets and trained support vector machines (SVMs) to classify events as SOM, PV, or Mixed using the vector of pyramidal activity alone at each timepoint during the event. Classification performance rose above chance levels more than 1 s before the IN event onset (Figure 4B; see STAR Methods), and then peaked 0.70 and 0.63 s after the onset for the two- and three-way classifiers respectively, indicating the time of maximal IN effect after the onset of an event (Figure 4B). Control analysis revealed classification was not related to differences in mean activity, as the mean vector across all trials was approximately parallel to the decision boundary in each dataset for each classification method (Figures S5C and S5D). Together, the classification results suggest that specific patterns of activity in the local PYR population precede PV and SOM events. Although the origin of that difference is unclear from our study, it could reflect changes in the activity of extrinsic sources of input or neuromodulatory tone.

Populations of neurons encode information with varying levels of sparseness. This continuum ranges from dense codes, where all neurons more equally contribute, to local codes where only a small proportion of neurons are active^33^. We hypothesized, based on previous results where optogenetic activation of SOM cells increases sparsity of sensory responses^14^, that PYR population activity would be sparser during SOM events than during PV events. To examine the sparsity of population activity, we examined the angle between the mean population vector during events to the unit vector, which would correspond to all neurons in the population being equally active. A greater angle between two population vectors suggests that the relative activity between neurons in the population is less similar^14,33,34^. Therefore, in comparing a population activity vector measured during IN events to the unit vector, we can determine the similarity between the actual pattern of activity and a dense code (the unit vector). We report these angles in terms of cosine similarity, where an angle of zero between the vectors would indicate identical ratios of activity between neurons, and result in a cosine similarity of one. We found that PYR activity during SOM events was sparser than during Mixed or PV events (p = 0.0015, p = 0.00049, Wilcoxon signed-rank), reflected by lower cosine similarity between the population vector and the unit vector, where all neurons would be equally active (Figure 4C). In tandem with increased sparseness, SOM neuron stimulation increases angle between mean population vectors in response to different stimuli, enhancing the separability of evoked population activity patterns^14^. We observed a similar effect in spontaneous activity, as the pairwise similarity between population activity vectors was lowest across SOM events than other IN event types (SOM/PV, p = 0.0049; SOM/Mixed, p = 0.0044; PV/Mixed, p = 0.09, Wilcoxon signed-rank; Figure 4D). This indicates that more population states were possible during events with both higher SOM neuron activity and lower PV neuron activity.

Our results indicate that pyramidal cells fired more independently when SOM activity was high and PV was low, reflected by increased sparseness and variability of PYR activity across SOM activity events. Prior work in V1 has shown that SOM neuron stimulation can modulate the coherence of population activity, lowering pairwise activity correlations between neurons^30^. Pairwise correlations between PYR neurons were indeed lower during SOM events compared to PV or Mixed events (SOM/PV, p = 0.020, SOM/Mixed, p = 0.032; PV/Mixed, p = 0.070, Wilcoxon signed-rank; Figure S5E), which suggests that natural bursts of activity in SOM neurons may also decorrelate local activity, like optogenetic stimulation of SOM neuron populations. Shared variability tends to be low-dimensional, where a large proportion of shared variability can be explained by a single latent variable^35^. To compare the dimensionality of shared variability within the PYR population during SOM, PV, and Mixed events, we used factor analysis to find axes in high dimensional space along which many neurons covary. The top five latent factors explained less shared variability across all periods of time during SOM events, indicating pyramidal neurons acted more independently during these times (Figure 3E; see Table S3 for statistics). Since these results could be explained either by our finding that mean activity during SOM events can occupy more possible states, or a reduction in covariance between neurons on shorter timescales during an event, we additionally took pairwise correlations of the mean-subtracted activity between neurons within each individual event, for only active neurons. Pyramidal neurons had lower correlations within SOM events than Mixed or PV events (SOM/PV, p = 0.015, SOM/Mixed, p = 0.032; PV/Mixed, p = 0.77, Wilcoxon signed-rank; Figure 4F), suggesting our results from factor analysis were explained by lower moment-to-moment fluctuations shared between neurons as well as the comparatively greater span of neural activity across SOM events.

## Discussion

Here we have demonstrated a simple and reliable method for distinguishing two red fluorophores *in vivo*, leveraging differences in their excitation spectra, allowing us to monitor the activity of two specific inhibitory cells along with the rest of the neural population, using a standard two-photon microscope with two detection channels. When used to identify cortical interneurons, any combination of two of the three main classes of INs allows the experimenter to comfortably approximate the remaining neurons as pyramidal, as they will comprise approximately 95% of the unlabeled population^4^. This enables analysis methods to parse how interactions between interneurons, which strongly connect to one another^13^, affect activity in the broader pyramidal population. When combined with experiments using optogenetic stimulation, the presentation of sensory stimuli, or involving goal-directed behavior, this will allow a more complete explanation of neural activity and its connection to behavior on individual trials.

Past research in sensory cortex has shown that IN activity tends to be highly coordinated, especially within cell type^17,25,27^. Our results in PPC, an association area that integrates multimodal sensory input^36–38^, demonstrate this may be a general feature in cortical circuits. By labeling the two IN subtypes that strongly inhibit different compartments of pyramidal cells^4^, we could isolate periods of time when coordinated IN activity occurred and characterize the cooccurring network activity state in the pyramidal population during PV and SOM events. This would not have been possible by labeling one IN at a time: pyramidal activity during Mixed events resembled PV events more closely than SOM events (Figures 4C-4F and S5E), even though SOM activity was also relatively high in Mixed events (Figure 3B). Our labeling also revealed pyramidal neurons that preferred either PV or SOM events (Figures 4A and S5A). These neurons were differently correlated with the rest of the local population, indicating that they perhaps have distinct functional roles. Furthermore, these neurons had strong correlations with one another, but not to neurons of opposite preference, suggesting they may form functional subnetworks (Figures S5A and S5B). In agreement with previous results^25^, pyramidal neurons were differentially involved in driving the relative IN activity of the population, as suggested by the ability of a linear decoder to predict the type of event from the preceding pyramidal activity (Figure 4B).

The information that can be encoded by populations of neurons is constrained by the correlation structure of their activity, by restricting the number of possible states a population can occupy^39^. This reduction in states due to the dependence between neurons can be conceptualized as lowering the dimensionality of population activity^40^. Across analyses at different time scales of the event types, a general pattern emerged where pyramidal neuron activity was more independent when SOM activity was high than when PV activity was high. This was reflected in more sparse activity during SOM events (Figure 4C) that could span a broader set of population states (Figure 4D). Factor analysis was better able to summarize the population with fewer dimensions across all PV events, and within events pairwise correlations were lower from moment-to-moment (Figures 4E and F). This is despite the fact that SOM neurons are more coordinated in their activity (Figure 2E; Table 1), which would indicate *a priori* that activity states identified by their mean activity would be more homogeneous.

Our findings of increased pyramidal independence with greater SOM neuron activity are corroborated by similar results from auditory and visual cortex in response to optogenetic stimulation^14,30^, as well as increased SOM activity from larger stimuli^41^. These studies have also noted the suppression of PV neurons corresponding to the population effects observed from elevated SOM activity^2,9,41^, and one model found the inhibition of PV by SOM to be essential for explaining the effect^2^. In our data from association cortex, we observed more independent activity only when mean SOM activity was relatively high and mean PV activity was relatively low. While far from conclusive, this suggests that there is a conserved microcircuit involving SOM and PV neurons that can influence the independence of neural activity in different cortical areas. One potential mechanism comes from the distinct modes of inhibition provided by SOM and PV neurons, where inhibition of distal dendrites from SOM neurons may reduce recurrent excitation received from surrounding pyramidal neurons, allowing them to act more independent of one another^41^.

Controlling the correlations between neurons or dimensionality of the population has broad implications for the neural code. Depending on their structure, correlations can negatively influence the resolution of stimulus encoding^39,42–44^, and can be modified by attention and learning to improve task performance^45,46^. However, the relationship between correlations and behavioral performance is inverted in PPC^47,48^, seeming to sacrifice the resolution of information for better signal propagation^44^. Given the similarities between IN activity and correlations from our data in PPC compared to sensory cortex, modulatory input to interneurons in PPC may be different relative to sensory areas to best drive population activity and behavior. Further efforts to understand the underlying circuitry will better allow us to make predictions about how the brain achieves population dynamics that serve the processing demands of an individual brain region.

## Limitations of Study

Our analysis in this paper is strictly correlational. We are not controlling input to the network with sensory stimuli, goal-directed behavior, or optogenetics. We additionally are only using the mean activity of interneurons to make conclusions about the functions of cell types. As shown by recent experiments, there is heterogeneity within cell types that will not be captured by the mean^49^. A more complete understanding of the function of these neuron classes will need to come from combining our approach with more controlled experiments such as perceptual decision-making tasks, which allow for looking at Ins based on their function as they correspond to task performance or stimulus features.

## Acknowledgments

We thank members of the Runyan lab for comments on the manuscript. We thank the GENIE project (Janelia) for making GCaMP sensors available for use. We thank the developers of Suite2P and Wavesurfer. This work was supported by the Andrew W. Mellon Predoctoral Fellowship, NIH Predoctoral Training Grant in Basic Neuroscience (T32 NS007433-24, NS007433-21), Pew Biomedical Scholars Program, the Searle Scholars Program, the Klingenstein-Simons Fellowship Award in Neuroscience, and NIH grants NIMH DP2MH122404, NINDS R01NS121913, NIH Supplement NIMH 3DP2MH122404-01S1.

## Author Contributions

C.A.R. and C.T.P. designed the experiments. C.D.B. conducted experiments. C.D.B. and C.T.P. processed data. C.T.P. performed analyses. C.A.R. and C.T.P. wrote the paper.

---

**Table S1.** Statistics for correlations in Figure 2.

|  | Pair Type | Mean Pairwise Correlation | std | n pairs |
| --- | --- | --- | --- | --- |
| 1 | PYR-PYR | 0.0184 | 0.0503 | 573661 |
| 2 | SOM-SOM | 0.0992 | 0.1279 | 1471 |
| 3 | PV-PV | 0.0673 | 0.1316 | 10021 |
| 4 | SOM-PV | 0.0429 | 0.0896 | 6720 |
| 5 | PYR-SOM | 0.0231 | 0.0630 | 53661 |
| 6 | PYR-PV | 0.0284 | 0.0637 | 135258 |
|  | Permutation Test p-value | Mean Difference | Hedges' g |  |
| 1 vs 2 | .0014 | -0.0831 | -1.6408 |  |
| 1 vs 3 | .0014 | -0.0490 | -0.9285 |  |
| 1 vs 4 | .0014 | -0.0255 | -0.5015 |  |
| 1 vs 5 | .0014 | -0.0047 | -0.0917 |  |
| 1 vs 6 | .0014 | -0.0094 | -0.1773 |  |
| 2 vs 3 | .0014 | 0.0341 | 0.2600 |  |
| 2 vs 4 | .0014 | 0.0575 | 0.5895 |  |
| 2 vs 5 | .0014 | 0.0784 | 1.1946 |  |
| 2 vs 6 | .0014 | 0.0737 | 1.1380 |  |
| 3 vs 4 | .0014 | 0.0234 | 0.2011 |  |
| 3 vs 5 | .0014 | 0.0443 | 0.5679 |  |
| 3 vs 6 | .0014 | 0.0396 | 0.5611 |  |
| 4 vs 5 | .0014 | 0.0208 | 0.3129 |  |
| 5 vs 6 | .0014 | -0.0047 | -0.0739 |  |

**Table S2:** Comparison of frequency and duration of events across event types.

| Frequency Across Datasets (Events Per Minute) | Event Type | Mean | Standard Deviation |  | Comparison (Wilcoxon Signed-Rank) | p-value |
| --- | --- | --- | --- | --- | --- | --- |
|  | SOM | 1.1982 | .4328 |  | SOM vs PV | .2661 |
|  | PV | .9945 | .2815 |  | SOM vs Mixed | .2661 |
|  | Mixed | .9421 | .3539 |  | PV vs Mixed | .7646 |
| Onset to Half-Max Duration of All Events (Seconds) |  |  |  |  | Comparison (Wilcoxon Rank-Sum) |  |
|  | SOM | 3.3026 | 1.9340 |  | SOM vs PV | .9953 |
|  | PV | 3.3726 | 2.0371 |  | SOM vs Mixed | .7229 |
|  | Mixed | 3.3726 | 2.0984 |  | PV vs Mixed | .7400 |
| Peak to Half-Max Duration of All Events (Seconds) |  |  |  |  | Comparison (Wilcoxon Rank-Sum) |  |
|  | SOM | 1.9965 | 1.3361 |  | SOM vs PV | .9138 |
|  | PV | 2.0493 | 1.4020 |  | SOM vs Mixed | .8297 |
|  | Mixed | 2.0507 | 1.4066 |  | PV vs Mixed | .7871 |

**Table S3.** Statistics and P-values for each comparison of factor analysis in Figure 4E using Wilcoxon rank-sum.

| Factor | Event Type | Mean | s.d. | 95% Confidence Interval |  |
| --- | --- | --- | --- | --- | --- |
| 1 | SOM | .0162 | .0093 | .0137 - .0186 | 518 |
|  | PV | .0231 | .0132 | .0197 - .0265 | 519 |
|  | Mixed | .0244 | .0126 | .0211 - .0276 | 520 |
| 2 | SOM | .0109 | .0070 | .0091 - .0127 | 521 |
|  | PV | .0154 | .0095 | .0130 - .0265 | 522 |
|  | Mixed | .0165 | .0089 | .0142 - .0188 | 523 |
| 3 | SOM | .0084 | .0059 | .0071 - .0096 |  |
|  | PV | .0110 | .0074 | .0091 - .0129 | 524 |
|  | Mixed | .0120 | .0065 | .0103 - .0188 | 525 |
| 4 | SOM | .0066 | .0030 | .0058 - .0074 |  |
|  | PV | .0087 | .0049 | .0074 - .0099 | 526 |
|  | Mixed | .0099 | .0056 | .0103 - .0114 | 527 |
| 5 | SOM | .0048 | .0022 | .0043 - .0054 |  |
|  | PV | .0069 | .0037 | .0059 - .0078 | 528 |
|  | Mixed | .0078 | .0043 | .0067 - .0089 | 529 |

| P – values | Factor 1 | Factor 2 | Factor 3 | Factor 4 | Factor 5 |
| --- | --- | --- | --- | --- | --- |
| SOM vs PV | .0033 | .0051 | .0983 | .0983 | .0033 |
| SOM vs Mixed | .00014 | < .0001 | .0012 | .0007 | <.0001 |
| PV vs Mixed | .4323 | .1493 | .1493 | .0983 | .0883 |

## Methods

### Lead contact

Further information and requests for resources and reagents should be directed to and will be fulfilled by the lead contact, Caroline A. Runyan.

### Materials availability

This study did not generate unique new reagents.

### Experimental Model and Subject Details

All procedures were approved by the University of Pittsburgh Institutional Animal Care and Use Committee. Homozygous SOM-Flp mice (Cat# 31629, Jackson Laboratory, USA) were crossed with homozygous PV-Cre mice (Cat# 17320, Jackson Laboratory, USA) obtained from, and all experiments were performed in the F1 generation, which expressed Flp in SOM+ neurons and Cre in PV + neurons. Mice were group housed in cages with between 2 and 4 mice. Adult (8–24 weeks) male and female mice were used for experiments (4 male, 3 female). Mice were housed on a reversed 12 hr light/dark cycle, and all experiments were performed in the dark (active) phase.

## Method Details

### Surgery

12-24 hours prior to surgery, mice were given a subcutaneous injection of dexamethasone (Covetrus, ME). Mice were anesthetized with isoflurane (4% for induction, 1-2% maintenance during surgery depending on breathing rate). Mice were mounted on a stereotaxic frame (David Kopf Instruments, CA). Subcutaneous injections of carprofen (Covetrus, ME) and dexamethasone were injected subcutaneously immediately prior to surgery for pain management and to reduce the inflammatory response. Ophthalmic ointment was applied to the eyes to prevent drying (Henry Schein, NY). A 4 x 4 mm craniotomy was made over PPC (centered at 2 mm posterior and 1.75 mm lateral to bregma) with a hand drill. A 1:1:2 mixture of three viruses was loaded into the glass injection pipet (1, Addgene #114471 pAAV-Ef1a-fDIO-mCherry, 2, Addgene #28306 pAAV-FLEX-tdTomato, and 3, Addgene #100837 pAAV.Syn.GCaMP6f.WPRE.SV40A). pAAV-Ef1a-fDIO mCherry was a gift from Karl Deisseroth (Addgene viral prep # 114471-AAV1; http://n2t.net/addgene:114471; RRID:Addgene_114471). pAAV-FLEX-tdTomato was a gift from Edward Boyden (Addgene viral prep # 28306-AAV1; http://n2t.net/addgene:28306; RRID:Addgene_28306). pAAV.Syn.GCaMP6f.WPRE.SV40 was a gift from Douglas Kim & GENIE Project (Addgene viral prep # 100837-AAV1; http://n2t.net/addgene:100837; RRID:Addgene_100837). A micromanipulator (QUAD, Sutter, CA) was used to target injections ∼250 μm under the dura at each site, where ∼60 nL virus was pressure-injected over 5-10 minutes. Pipets were not removed until 5 minutes post-injection to prevent backflow. Dental cement (Parkell, NY) sealed a glass coverslip (4mm) over a drop of Kwik Sil (World Precision Instruments, FL) over the craniotomy. Using dental cement, a one-sided titanium headplate was attached to the right hemisphere of the skull. After mice had recovered from the anesthesia, they were returned to their home cages, and received oral carprofen tablets (Fisher Scientific, MA) for 3 days post-surgery.

### Control Surgery

We performed our craniotomy surgery with the methods described above, except there were physically separated injections of the mCherry and tdTomato viral constructs at neighboring sites in each animal (six total injections). The mCherry virus was injected medially, while the tdTomato virus was injected laterally, with 1 mm separation in the mediolateral axis. The dilution of the virus was the same in all control and experimental animals, however one control animal received injections containing only the fluorophore viruses (diluted 1:10 with sterile 1X PBS) while the other received control injections that also included the GCaMP virus (2:1:1 of GCaMP/red-expressing virus/PBS).

### Two-photon microscope

Images were acquired using a resonant scanning two-photon microscope (Ultima Investigator, Bruker, WI) at a 30 Hz frame rate and 512 x 512 pixel resolution through a 16x water immersion lens. PPC was imaged at a depth between 150 and 250 μm, at the level cortical layer 2/3. The angle of the objective was matched to the angle of the window. Excitation light was provided by both a tunable femtosecond infrared source (780-1100 nm) and a fixed 1045 nm wavelength laser (Insight X3, Spectra-Physics, CA). Tunable and fixed wavelength beams were combined with a dichroic mirror (ZT1040dcrb-UF3, Chroma, VT) before being routed to the microscope’s galvanometers. Note that because of this optics configuration, imaging cannot be performed at tunable wavelengths immediately surrounding 1045 nm. Green and red wavelengths were separated through a 565 nm lowpass filter before passing through bandpass filters (Chroma, ET525/70 and ET 595/50, VT). PrairieView software (vX5.5 Bruker, WI) was used to control the microscope.

### Behavioral Monitoring

Running velocity was monitored on pitch, roll, and yaw axes using two optical sensors (ADNS-98000, Tindie, CA) held adjacent to the spherical treadmill. A microcontroller (Teensy, 3.1, Adafruit, NY) communicated with the sensors, demixing their inputs to produce one output channel per rotational axis using custom code. Outputs controlling the galvanometers were synchronized with running velocity using a digital oscilloscope (DIGIDATA 155080, pClamp 11, Molecular Devices). Instantaneous velocity was calculated using the distance formula, using pitch and roll velocity values as inputs.

### Image Acquisition

Fields of view were selected based on GCaMP expression and red fluorophore expression. mCherry vs tdTomato expression could be quickly assessed by comparing images collected at 780 nm and 1045 nm excitation, as both fluorophores were visible at 1045 nm, but only mCherry was visible at 780 nm. Multiple imaging sessions were acquired from each cranial window, varying depths and locations rostro-caudally and medio-laterally. The wavelength series to distinguish mCherry and tdTomato fluorescence was performed first. Images were collected at excitation wavelengths from 780 nm to 1100 nm in increments of 20 nm each averaged over 16 frames, excluding 1020 and 1060 nm wavelengths. The 1040 nm image was acquired with the fixed-wavelength beam from the Insight laser. At each wavelength, images were taken at multiple powers (0 to 275 mW). Powers were chosen for each dataset post-imaging as to avoid ones that were collected at too high or too low a power. The laser was then tuned to 920 nm for functional imaging of GCaMP fluorescence. Images were collected while mice ran voluntarily on an air-suspended spherical treadmill^17^. Functional imaging sessions were approximately 30 minutes in duration. All images were collected at a 30 Hz frame rate.

### Image Processing

The images from the wavelength series and functional imaging were processed separately in Suite2p (v0.10.3) in Python with Cellpose (v0.7.3), along with 1000 frames of images collected during the functional imaging phase of the same imaging session, to allow for registration between the wavelength series and the functional imaging. Motion correction was performed on the functional images collected at 920 nm and on the wavelength series, only the functional imaging was used for ROI extraction. We then manually selected all red cells. Custom MATLAB code was used to load the two registered series of images, the wavelength series and functional imaging. Any discrepancy in registration between these two datasets was corrected by adding the peak lag of the cross correlation between the wavelength series and functional data in the X and Y directions to the coordinates of the ROIs. This correction was manually verified by plotting the circumference of the ROIs over the mean images from each image registration and confirming that the location of the ROIs relative to the cells were identical.

To further verify that each cell contained a red fluorophore, our MATLAB code was then used to plot outlines of the ROIs from the functional imaging. First, to rule out potentially ambiguous red neurons, we plotted these ROIs onto max projection of the wavelength series, as well as mean images taken from the 780-820nm range and the 960-1045nm range to best show mCherry or tdTomato containing cells respectively. Any ROI containing out of focus red expression or that contained overlapping red neuropil or a closely neighboring soma from another neuron was excluded from analysis entirely.

Having chosen which neurons contained red fluorophores, we generated vectors for each of the red fluorophore containing cells, where each dimension was the mean intensity of the pixels of each cell’s ROI at a given wavelength. Based on our results revealing the best wavelength combinations for separating tdTomato and mCherry from Figure 1E, we used the combination in the top 25 that has the least variability across datasets (see Figure 1E, *mCherry and tdTomato ROI classification*), but all combinations in the top 25 produce almost identical results after exclusion of ROIs below the silhouette score of 0.7. For our data we used the wavelengths corresponding to combination 5 in Figure 1E, consisting of the wavelengths: 780, 800, 820, 940, and 960 nm.

For the control data (Figure 1), since there was no functional data to generate ROIs in suite2p, ROIs were manually drawn. One of our two control mice had a GCaMP construct in each control injection, and there was no measured difference in clustering between control mice with and without GCaMP co-expression.

### mCherry and tdTomato ROI Classification

Vectors containing the mean intensity of all pixels for each ROI at each wavelength (see *Image Processing*) were run through the k-means clustering algorithm function in MATLAB. This algorithm uses a squared Euclidian distance metric for optimization and the k-means++ algorithm for cluster center initialization. Silhouette scores were calculated for each neuron using the following formula:

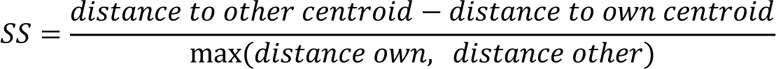

The k-means clustering algorithm was applied to every possible combination of imaging wavelengths (n=32752 combinations) described above on each of the 6 datasets from our control animals. We used the mean of the silhouette scores for all neurons in each iteration of this process as a metric of cluster separability. For all subsequent analyses, we used a specific wavelength combination that performed well across all control datasets, to maximize the stability of classification across datasets (780, 800, 820, 940, 960 nm). To determine a threshold by which cells would be excluded, we examined the distribution of silhouette scores of misclassified cells from all the iterations of clustering on control data. Note that these included ‘suboptimal’ combinations of wavelengths that do not distinguish mCherry and tdTomato as well. The silhouette scores from the bottom 10% of these iterations were excluded, as quantified by number of cells misclassified. This was done to remove datasets that were so noisy as to not be representative of what cluster separability would be like in intermingled data. We took two standard deviations above the mean of this distribution as a threshold for exclusion (0.7 SS). Any neuron in our experimental data that had a silhouette score lower than this value was excluded from analysis entirely. The cluster that included ROIs with strong fluorescence at 780 nm was identified as the mCherry/SOM cluster.

### GCaMP Fluorescence Processing

Images of GCaMP fluorescence collected with 920 nm excitation were processed in a separate suite2p instance (v 0.10.3, Cellpose 0.7.3). Raw imaging movies were processed to correct for motion, identify cell bodies, and extract soma and neuropil fluorescence. Suite2p generated regions of interests (ROIs) for each cell based on anatomical features and correlations between pixels across time during functional imaging. ROIs were manually verified to be cell bodies, as opposed to dendritic processed, based on their morphology. The fluorescence of the neuropil surrounding the cell body was subtracted off from the fluorescence of the cell body, weighted by a factor of 0.7. The resulting signal was used for extracting a dF/F and deconvolved signal for each cell. ROIs classified as mCherry+ were considered SOM neurons, tdTomato+ were considered PV neurons, and GCaMP+ neurons that had no red fluorescence were considered putative pyramidal (PYR) neurons, these neurons comprise >90% of cortical neurons that do not express PV or SOM^4^.

### dF/F and Deconvolution

Once the neuropil corrected fluorescence was obtained for each animal (F), dF/F was calculated for each neuron (F-Fbaseline)/Fbaseline for each frame. Fbaseline was the eighth percentile of the 900 F values surrounding that frame for a given neuron (∼15 s in each direction, 30 s total). dF/F timeseries were then deconvolved to estimate the relative spike rate in each imaging frame using the OASIS toolbox. We used the AR1 FOOPSI algorithm and allowed the toolbox to optimize the convolution, kernel, baseline fluorescence, and noise distribution. Deconvolved spike rates were thresholded by setting values below 0.05 to 0.

### Pairwise correlations

Partial Pearson correlations were computed between all pairs of neurons for the results in Figure 2, for all combinations of cell types: Pyr-Pyr, SOM-SOM, PV-PV, Pyr-SOM, Pyr-PV, SOM-PV. Activity was boxcar smoothed in time (300 ms), and the partial Pearson correlation between the activity of the neurons was computed, discounting the effect velocity in the pitch, roll, and yaw directions separately (Matlab function ‘partialcorr’). This procedure was the same for Figure S5A and F, except pairwise correlations were only calculated on the concatenated activity of non-event periods in S5A and the concatenated activity from different events in S5F. For pairwise correlations within trials in Figure 4F, all pairwise correlations were calculated for each event, again discounting the effect of running. The effects of mean activity were discounted by subtracting the mean activity for each event before making the correlation.

### Identification of PV and SOM population events

Coordinated PV, SOM, and Mixed events were identified for each imaging session as follows. The activity of individual PV and SOM neurons over the entire session was z-scored, and then each neurons activity was averaged together across cell type and box-car smoothed with a 300 ms window. Peaks in this population activity signal were identified from this average signal (MATLAB’s function ‘findpeaks’, MinPeakHeight: 1 standard deviation, MinPeakProminence: 2 standard deviations, MinPeakWidth: 10 ms). Peaks in each interneuron population were considered “Mixed” if they corresponded to elevated activity in the other interneuron population (> 1/2 of the normalized activity of the population signal where the peak was identified). The onset of the events was defined by the point at which the prominence was closest to zero in the period of time defined by half of the event width prior to the first half-max (see Figure S3A for event definitions). We additionally tried using parameters of half these values, but we did not observe differences in our results.

### Event Type Classification

We trained a support vector machine (SVM) to classify whether an event was SOM or PV based on mean activity across all events at time bins spanning from 3.3 seconds before the onset of the event until 1.66 seconds after that point. Bins were averaged over 300 ms. Training was done on n-matched numbers of events, and performance was calculated based on 10-fold cross-validation. An identical procedure was done to discriminate SOM, PV, and Mixed events, except a one-vs-one SVM was used. Statistical significance was calculated based on a permutation test done between 100 control classification accuracies using shuffled event type labels and our classification result for each time point.

### Population Vector Cosine Similarity

The cosine similarity between population vectors was calculated with the following formula:

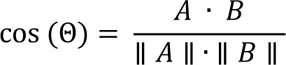

Where Θ is the angle between the population vectors A and B, and cos(Θ) is the cosine similarity. Population vectors were taken from the mean activity two seconds after the peak of the event. For calculating sparseness, we took the cosine similarity between each population vector and the unit vector, where each dimension had a value of 1. For calculating the cosine similarity of vectors within each event type, we took the cosine similarity between all pairwise combinations of vectors for a given event type.

### Factor Analysis

Factor analysis was used to measure the shared variability between neurons^50,51^ using the deconvolved responses of a randomly selected subset of 75 PYR neurons within each imaging session. Factor analysis was trained on a matrix containing the deconvolved activity of all neurons at all timepoints over the event to collect 5 latent factors for SOM, PV, and Mixed events. Deconvolved activity was square root normalized to avoid biasing towards highly active neurons^50^. Training was done using 10-fold cross validation. The eigenvalue for each latent factor corresponds to the PYR population’s shared variability in that dimension. This procedure was repeated 5 times for each dataset.

### Histology

After all imaging sessions had been acquired, each mouse was transcardially perfused with saline and then 4% paraformaldehyde. The brain was extracted, cryoprotected, embedded, frozen, and sliced. Once slide mounted, we stained brains with DAPI to be able to identify structure. We used anatomical structure to verify the locations of our injections in PPC.

## QUANTIFICATION AND STATISTICAL ANALYSIS

All pairwise comparisons across datasets were done with paired or unpaired non-parametric tests (Wilcoxon signed rank and Wilcoxon rank-sum). Comparisons performed directly on deconvolved data and also on classification results were done with unpaired permutation (i.e. randomization) tests with 10,000 iterations for deconvolved, where p<.0001 indicates the highest significance for the number of runs performed. 100 iterations were performed to determine classification performance significantly better than chance from classification performed on shuffled labels. Permutation tests and non-parametric tests were run to test differences in means.

---

**Figure S1.**
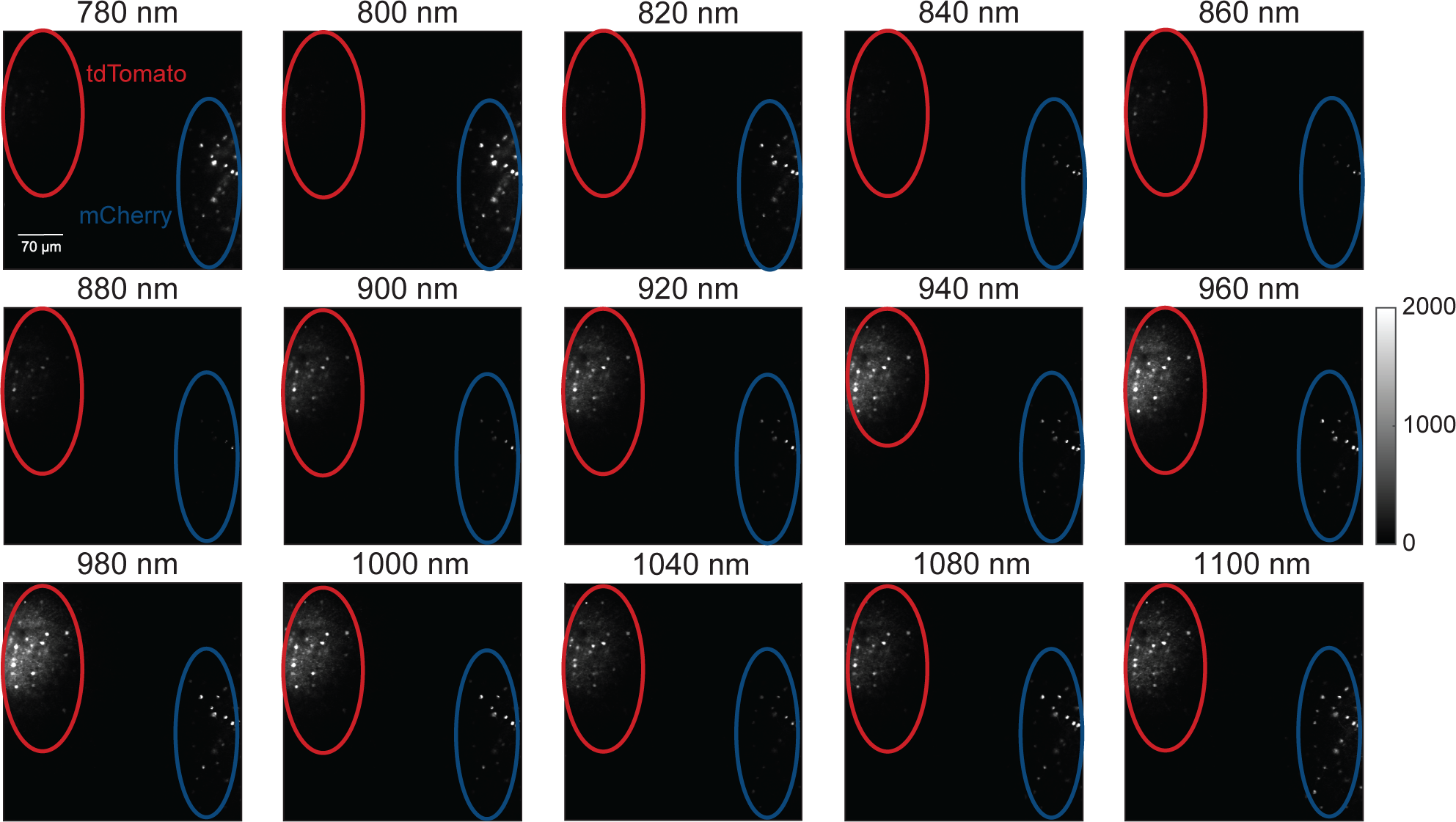
mCherry and tdTomato sites have visibly distinct excitation spectra. Example wavelength series in a control mouse with spatially segregated injection sites, demonstrating visible differences in excitation spectra between the two fluorophores. Circles indicate the location of the injection site, the red indicating tdTomato and the blue indicating mCherry. The color mapping in the images to the raw fluorescence intensity is the same across all pictures, the colorbar applies to all panels. Scalebar is 70μm and applies to all panels.

**Figure S2.**
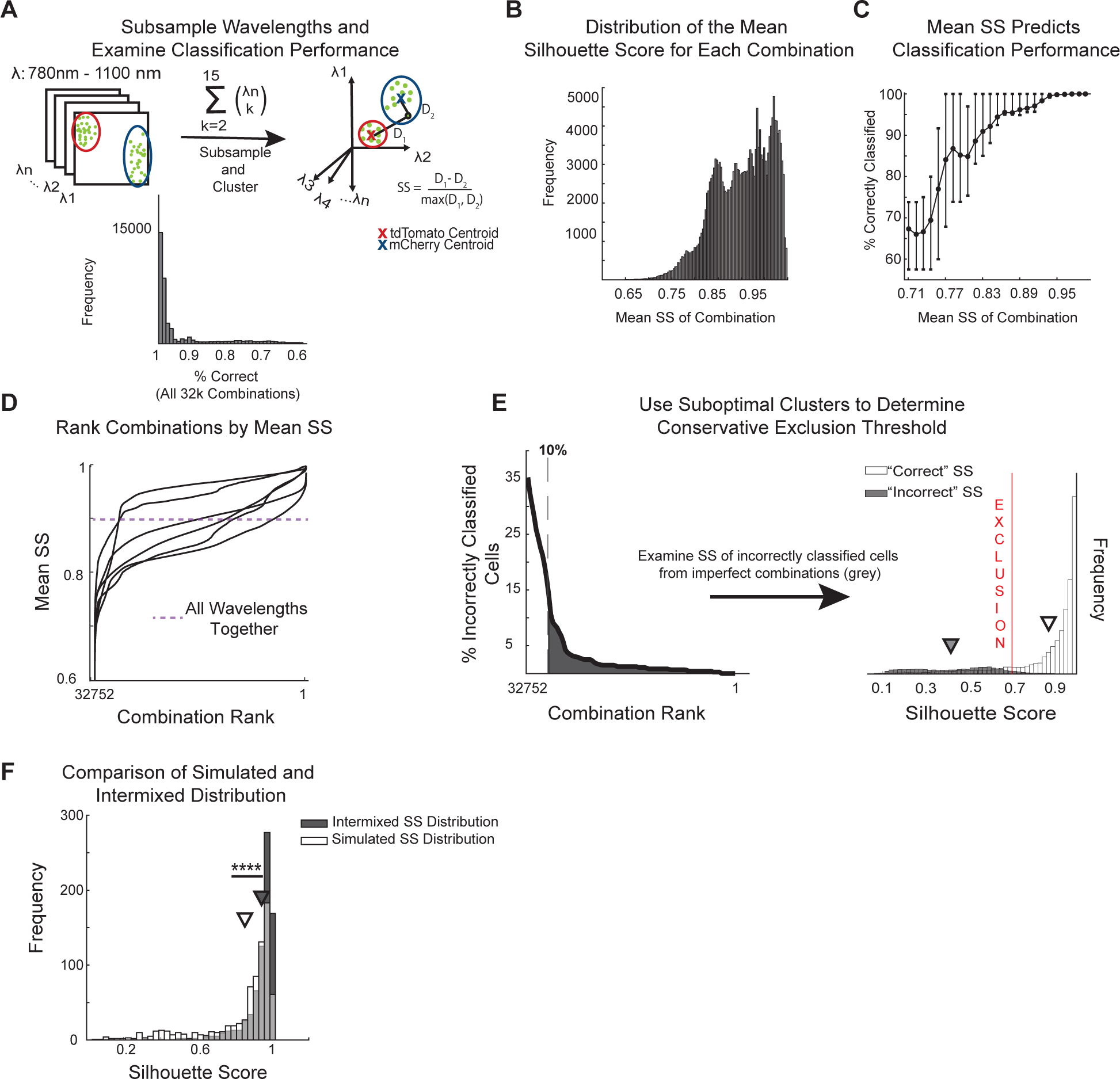
Determining optimal wavelength combination for clustering and a threshold for ROI exclusion. (A) (Top) Schematic describing subsampling procedure to find the combinations of wavelengths that resulted in the most separable clustering between mCherry and TdTomato ROIs. Clustering was attempted using all possible combinations of the 15 wavelengths and a silhouette score was computed for each neuron using each wavelength combination. (Bottom) Most wavelength combinations allowed accurate classification of fluorophore identity. (B) Distribution of the mean silhouette scores from clustering on each possible combination of the 15 wavelengths. (C) Relationship between the mean silhouette score of all cells after clustering and the classification performance, showing that high silhouette scores indicate reliable classification performance. Error bars represent the interquartile range of classification performance within each bin. (D) Combinations of wavelengths were ranked by their mean silhouette score. Each trace represents one dataset. The dashed purple line indicates the mean silhouette score for clustering using all wavelengths, demonstrating the improvement in separability when using the highest performing wavelength combinations. (E) (Left) To determine a threshold for excluding incorrectly classified cells, we examined the distribution of silhouette scores from incorrectly classified neurons using less informative wavelengths for clustering. We only drew from incorrectly classified cells that were obtained from the best 90% of ranked combinations, as below that clustering performance was noisier than could possibly be expected from real data. (Right) We defined a conservative threshold that would exclude 98% of misclassified neurons, even in conditions much noisier than in our control experiments. The distribution of “Correct” silhouette scores is subsampled so that the “Incorrect” silhouette scores are still visible. (F) Comparison of the distribution of silhouette scores obtained through using suboptimal wavelengths in the control data (white bars) versus those obtained experimentally with intermixed fluorophores (black bars). Simulated noisy data had worse clustering performance than experimentally obtained data, meaning that exclusion criteria were determined from a conservative estimate of noise in the data. Experimental data is n-matched to the simulated data to show relative differences in the distribution. For this panel, **** indicates p < 0.0001 on the Wilcoxon rank-sum test comparing the means of the two distributions.

**Figure S3.**
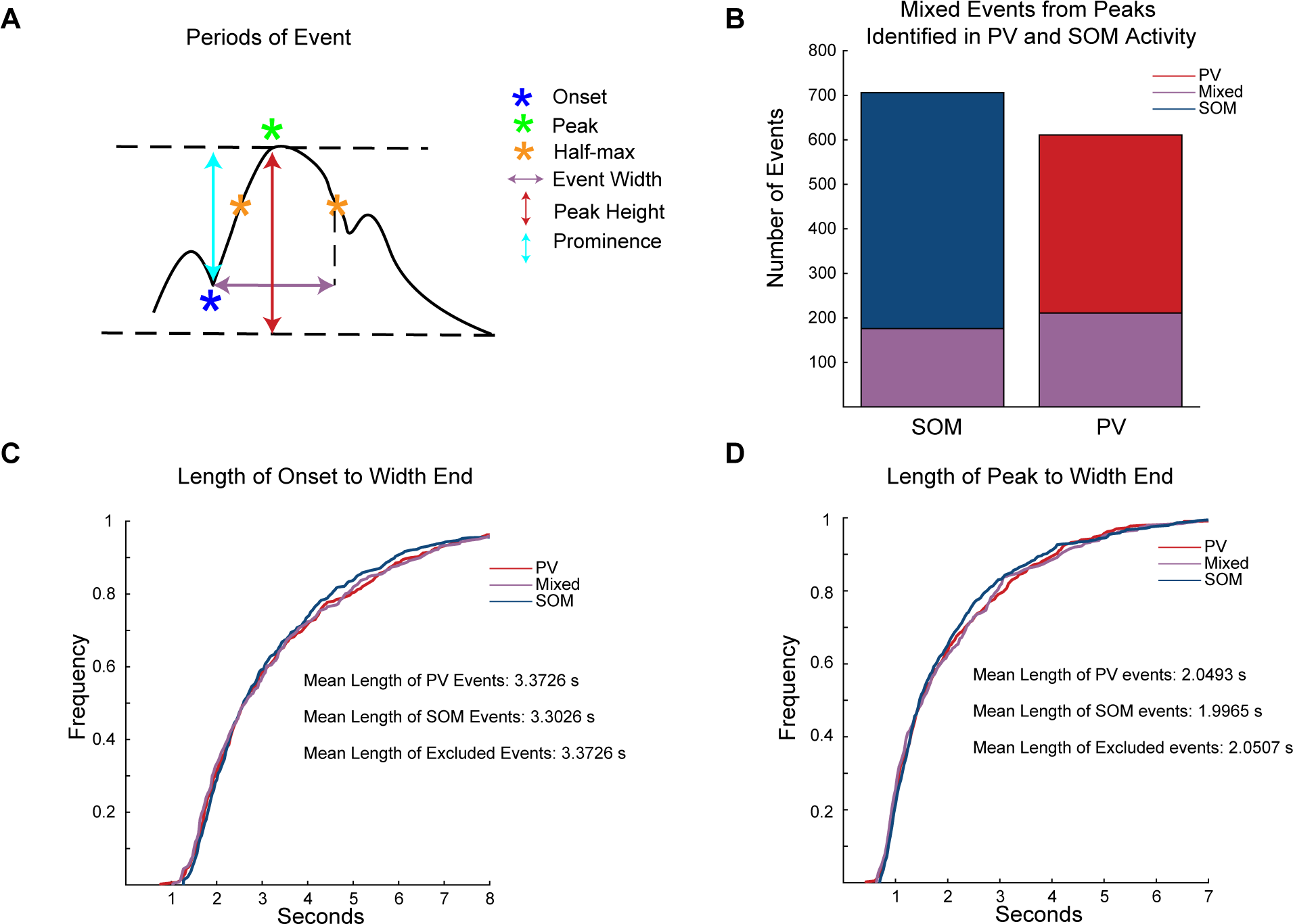
Description of event epochs and distribution of time periods across event types. (A) Schematic depicting how events were defined. To be considered an event, the peak had to be at >1 s.d. of the interneuron’s activity across the recording session. The prominence had to be >2 s.d. (B) Proportion of Mixed events that were taken from events initially identified by our peak finding algorithm from PV or SOM mean activity. (C) Distribution of the event durations from the onsets of events until the time of second half max. The events defined by our parameters were of similar length for each cell type. (D) The length of time from event peak to second half max was also similar for each cell type.

**Figure S4.**
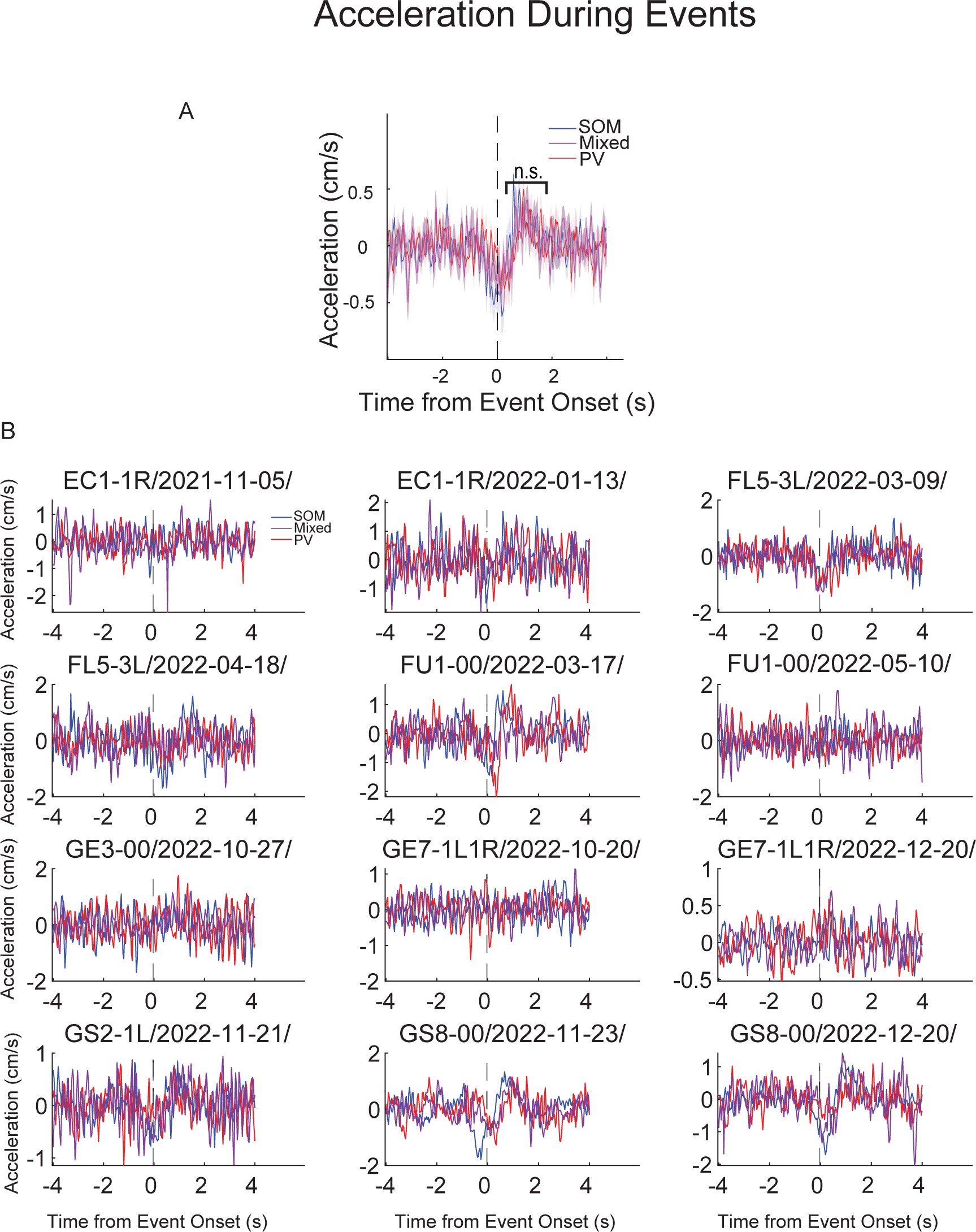
Differences in mouse running acceleration aligned to event onset were not observed across all datasets. (A) Instantaneous acceleration of the animal on the spherical treadmill. On average, acceleration slightly decreased for events of each cell type and then slightly increased. There was no significant difference between event types at the peak of the acceleration trace, indicated by “n.s.”. (B) Acceleration data from each dataset individually. The animal’s identifier and date of the recording session are indicated in the title of each panel. No consistent pattern was observed in all datasets.

**Figure S5.**
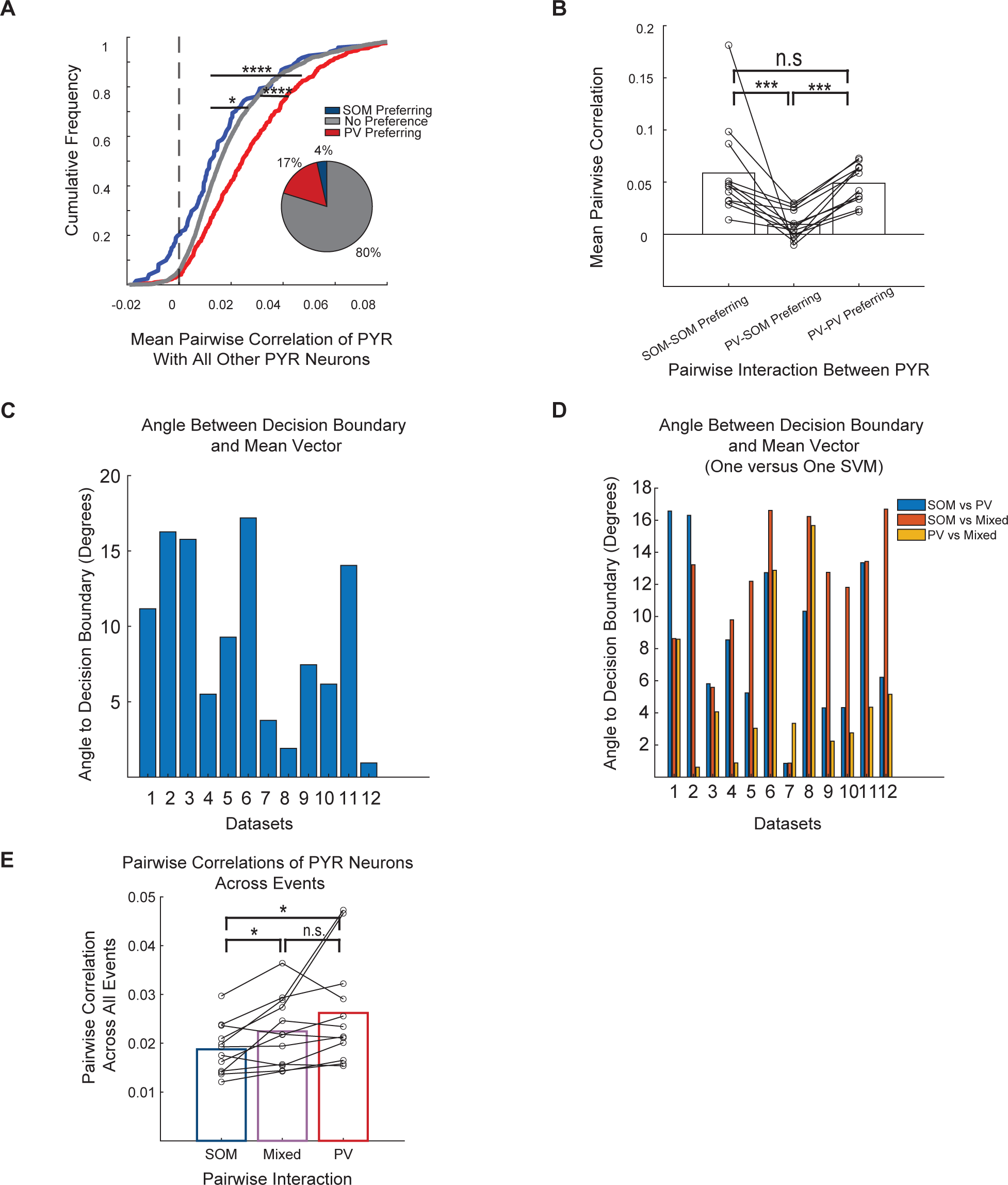
Distinct patterns of pyramidal activity across IN events. (A) Cumulative distributions of pairwise correlations between PYR neurons. Correlations were calculated across all timepoints that were not during an IN event. Preference was determined based on if the deconvolved activity events one of type was significantly different than for the other type in a permutation test (p<0.05). (B) Mean pairwise correlation between neurons that prefer the same event versus between those that prefer opposite events. As in (A), correlations were calculated on the concatenated activity during all timepoints that were not during an event. (C-D) Angle of the decision boundary to the mean PYR population activity vector across all SOM, PV, and Mixed events. An angle close to 90 would indicate that differences in mean activity were used by the decoder to classify event types. (C) Is for the binary classification between SOM and PV and D) is for the three-way classification using the one versus one SVM. (E) Average of the pairwise correlations between all pyramidal neurons across all timepoints for each event type. This is different from Figure 4F, where correlations were calculated for each event. For all panels, *, p<0.05; **, p<0.01; ***, p<0.001; ****, p<0.0001.

